# The evolutionary pathways for local adaptation in mountain hares

**DOI:** 10.1101/2021.09.16.460230

**Authors:** Iwona Giska, João Pimenta, Liliana Farelo, Pierre Boursot, Klaus Hackländer, Hannes Jenny, Neil Reid, W. Ian Montgomery, Paulo A. Prodöhl, Paulo C. Alves, José Melo-Ferreira

**Author notes:** **Corresponding author:** José Melo-Ferreira, CIBIO, Centro de Investigação em Biodiversidade e Recursos Genéticos, InBIO Laboratório Associado, Universidade do Porto, Campus de Vairão, 4485-661 Vairão, Portugal.

## Abstract

Understanding the evolution of local adaptations is a central aim of evolutionary biology and key for the identification of unique populations and lineages of conservation relevance. By combining RAD sequencing and whole-genome sequencing, we identify genetic signatures of local adaptation in mountain hares (*Lepus timidus*) from isolated and distinctive habitats of its wide distribution: Ireland, the Alps and Fennoscandia. Demographic modelling suggested that the split of these mountain hares occurred around 20 thousand years ago, providing the opportunity to study adaptive evolution over a short timescale. Using genome-wide scans, we identified signatures of extreme differentiation among hares from distinct geographic areas that overlap with area-specific selective sweeps, suggesting targets for local adaptation. Several identified candidate genes are associated with traits related to the uniqueness of the different environments inhabited by the three groups of mountain hares, including coat colour, ability to live at high altitudes and variation in body size. In Irish mountain hares, a variant of *ASIP*, a gene previously implicated in introgression-driven winter coat colour variation in mountain and snowshoe hares (*L. americanus*), may underlie brown winter coats, reinforcing the repeated nature of evolution at *ASIP* moulding adaptive seasonal colouration. Comparative genomic analyses across several hare species suggested that mountain hares’ adaptive variants appear predominantly species-specific. However, using coalescent simulations we also show instances where the candidate adaptive variants have been introduced via introgressive hybridization. Our work shows that standing adaptive variation, including that introgressed from other species, was a crucial component of the post-glacial dynamics of species.

## Introduction

During their evolution, species face environmental changes that challenge their distributions, potentially leading to local extinctions, range shifts or new adaptations (Nogués-Bravo et al., 2018). These changes in local environmental conditions and the colonization or recolonization of habitats with distinct selective pressures are potential triggers of divergent local adaptation. Identifying instances of adaptive evolution is a central aim in evolutionary biology, and also provides important insights to delineate conservation strategies aiming at preserving adaptive uniqueness and evolutionary potential (Funk, McKay, Hohenlohe, & Allendorf, 2012; Mills et al., 2018; Teixeira & Huber, 2021). For example, the existence of locally adapted populations across the range of widely distributed species may justify delineating guided conservation actions to preserve adaptive genetic variation. The advent of high throughput sequencing provides exceptional tools to detect signatures of adaptation along the genome, which can be linked to the morphological, physiological, behavioural, and ecological uniqueness of populations, and ultimately to fitness (Savolainen, Lascoux, & Merila, 2013; Hendricks, Schweizer, & Wayne, 2019). However, establishing a relationship between genetic variation and its adaptive underpinnings is challenging because it implies differential fitness in alternative habitats, which is not trivial to test. Scanning the genome for extraordinary differentiation between recently diverged populations, that is incompatible with neutral demographic expectations, can be used as an important first step towards identifying candidate genomic regions, genes and evolutionary pathways that underlie trait differences between locally adapted populations (Hoban et al., 2016).

The mountain hare (*Lepus timidus*) is an arctic-alpine species currently widely distributed in the northern Palearctic, from Fennoscandia to East Russia, with some isolated populations in Ireland, the Alps, Scotland, Faroe Islands and Poland, for example (Angerbjorn, 2018). Pleistocene fossil records show that the species was distributed further south at the Last Glacial Maximum (LGM), as in the Iberian Peninsula or Southern France (Altuna, 1970; Lopez-Martinez, 1980). Phylogeographic analyses of mitochondrial DNA and microsatellite variation (Hamill, Doyle, & Duke, 2006; Melo-Ferreira et al., 2007) as well as ancient DNA (Smith et al., 2017) have shown that the fragmented populations of the mountain hare share substantial variation, suggesting that they have diverged recently, potentially from a large range south of the ice rim. Thus, the current distribution of mountain hares is likely to have resulted from population fragmentation and colonization of deglaciated areas. The evolutionary history of the species, however, is far from established. Contemporary isolated mountain hare populations in Europe inhabit diverse biogeographical regions and habitats, such as the insular Atlantic habitats of Ireland, the alpine environment of the Alps, or the boreal forests and tundra of Scandinavia. Mountain hares from these isolated areas are characterised by distinct traits that may have arisen from recent local adaptation. These include the distinctive brown winter coat colour of the Irish mountain hare (instead of the typical summer-brown to winter-white pelage colour change), the smaller body size of the Alpine mountain hare and its ability to inhabit high altitudes, or the larger body size of Scandinavian mountain hare. These differences have supported their taxonomic classification into distinct subspecies, *L. t. hibernicus* in Ireland, *L. t. varronis* in the Alps, or *L. t. timidus and L. t. sylvaticus* in Fennoscandia (Angerbjörn & Flux, 1995). Furthermore, the range shifts of mountain hares since the LGM have led to transient or persistent contacts with other species of hares. These contacts have resulted in hybridization with non-negligible levels of introgression between mountain hares and other hare species (Melo-Ferreira, Alves, Freitas, Ferrand, & Boursot, 2009; Seixas, Boursot, & Melo-Ferreira, 2018; Giska et al., 2019). Often, these introgression events resulted from ancient species interactions, such as between the currently allopatric mountain and the Iberian hare (*L. granatensis*), which may have been driven by natural selection in many instances (Melo-Ferreira et al., 2012; Seixas et al., 2018; Giska et al., 2019). The mountain hare is therefore a privileged model to assess genomic signatures of selection resulting in local adaptation and to understand the evolutionary origin of adaptive variants.

Here, we analyse restriction site-associated DNA and whole-genome sequencing data from three isolated populations of mountain hares. We perform genome scans of differentiation and selection to identify candidates for local adaptation and genetic uniqueness. Using genomic data across several species of hares, we dissect the evolutionary trajectories underlying the candidate adaptive variation. We then discuss the relevance of our findings in the context of conservation strategies for the species.

## Methods

### Sampling

Mountain hares from Ireland, the Alps, and Fennoscandia are included in this study (Figure 1). These three regions represent disjoint ranges of species’ distribution, have very distinct environmental and climatic envelopes and harbour mountain hares with distinctive morphological features that could have resulted from local adaptation. Irish mountain hares were sampled in five counties (Mayo, Cavan, Limerick, Laois, Waterford), Alpine hares originated from France, Italy, Switzerland as well as Austria, and Fennoscandian hare samples derived from Sweden, Norway and Finland (see sampling details in Table S1). Hereafter we will use “population” to designate each sampled geographic region: Ireland, Alps and Fennoscandia.

**Figure 1.**
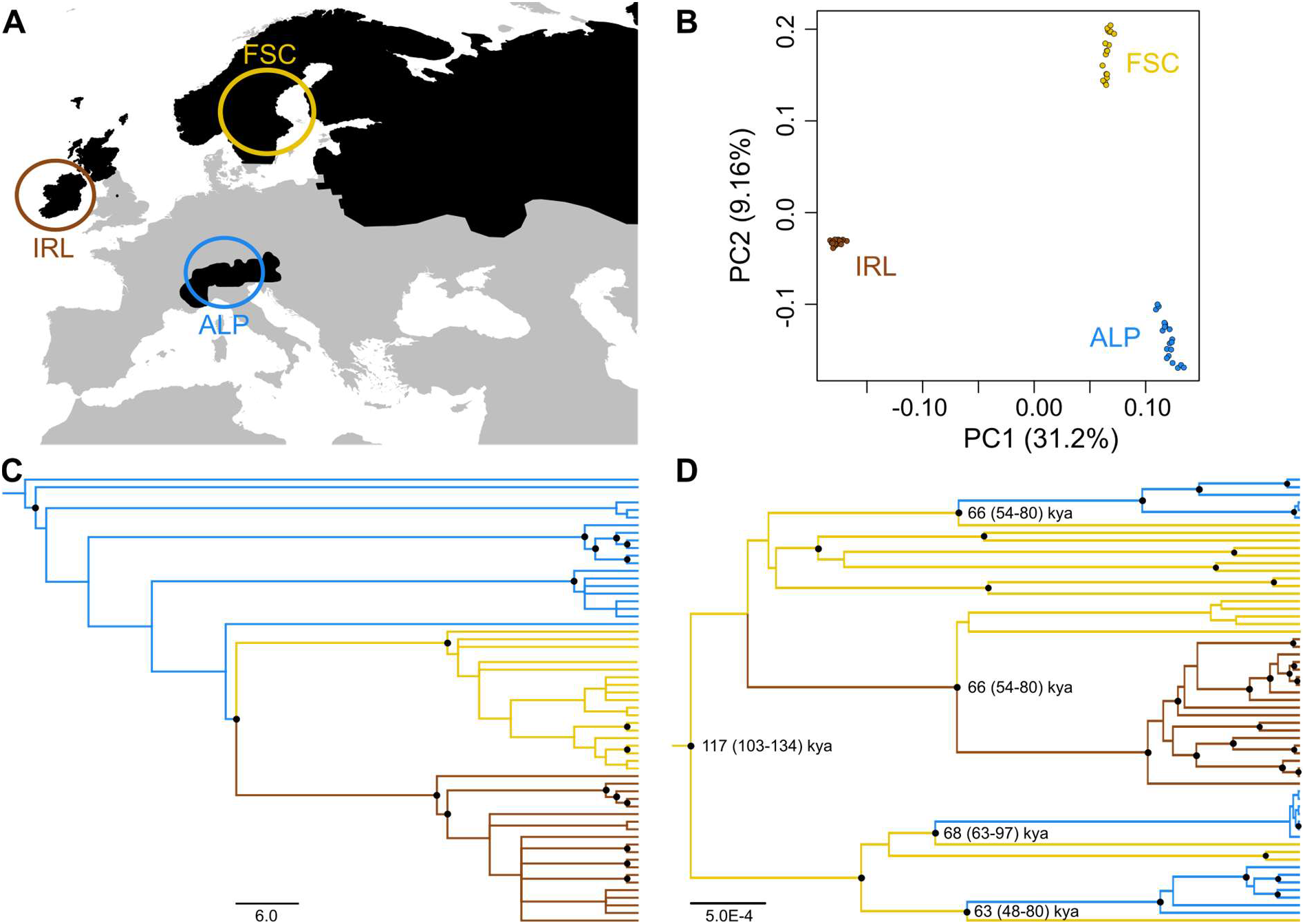
Mountain hare sampling and genetic relationships. (A) Distribution of the mountain hare in western Europe (black), according to Mitchell-Jones et al. (1999), and sampled regions: Ireland - IRL, Alps - ALP, Fennoscandia - FSC. (B) Principal component analysis (PCA) based on whole-genome data, inferred using 125,196 polymorphic sites, sampled at least 20 kb apart along the genome. (C) Neighbour-joining tree based on individual pairwise genetic distances estimated from 128,019 SNPs derived from whole-genome datasets. Nodes with bootstrap support values ≥ 0.90 are shown as black circles. (D) mtDNA phylogeny. Ages of key nodes are displayed (mean and 95% HDI) and nodes with posterior probability ≥ 0.90 are shown as black circles.

### RAD sequencing

Double-digest restriction site-associated DNA sequencing (RAD-seq) libraries were prepared for a total of 62 mountain hares from Ireland (N = 20), Fennoscandia (N = 26), and the Alps (N = 16) (Figure 1A, Table S1), following a protocol adapted from Peterson et al. (2012). Genomic DNA was extracted using the EasySpin Genomic DNA tissue kit (Citomed) and quantified with Qubit® 2.0 fluorometer (Invitrogen). For each individual, up to 400 ng of genomic DNA was digested with the SbfI-HF and MspI restriction enzymes (New England BioLabs) for 3h at 37°C, purified, quantified and ligated to adapters with compatible overhangs (P1-SbfI with a 5-7bp inline barcode, and P2-MspI). Individual DNA samples were then combined in pools based on (i) DNA intactness, assessed with gel electrophoresis before digestion and (ii) DNA concentration measured with Qubit after digestion but before ligation (the same amount of total DNA per sample was pooled). After electrophoresis on a 2.5% agarose gel, fragments between 450 and 700 bp were excised and purified with the MinElute Gel Extraction kit (Qiagen). Indexed Illumina sequencing adapters were added by PCR enrichment of ligated fragments with Phusion High-Fidelity DNA Polymerase using 10-12 cycles (Thermo Fisher Scientific). Amplified libraries were analysed on a 2200 TapeStation (Agilent Technologies) to confirm the size range of the recovered fragments. After quantification by qPCR, libraries were pooled in equimolar ratios and sequenced as part of two lanes of an Illumina HiSeq 2500 platform (paired-end, 150 bp reads) at Novogene (Novogene Corporation Ltd.).

The raw RAD-seq reads were demultiplexed based on the unique barcode-index combination and filtered using the program *process_radtags* included in the Stacks 2.52 pipeline (Catchen, Hohenlohe, Bassham, Amores, & Cresko, 2013) with the following filters: -c -q -r --retain_header --adapter_mm 1 -w 0.1 -s 20 -t 143 --len_limit 143. Reads were mapped onto a hare pseudo-reference, originally generated through iterative mapping of reads from three hare species onto the rabbit genome (Seixas et al., 2018; Seixas, Boursot, & Melo-Ferreira, 2019), which allowed retrieving the ancestral state and polarizing variants for downstream analyses. We used BWA-MEM with default parameters (Li & Durbin, 2009) to map the reads, which were locally realigned around indels using the RealignerTargetCreator and IndelRealigner functions from GATK v.3.8.1 (McKenna et al., 2010). Then we ran the program *ref_map*.*pl* from Stacks for loci assembly, SNP calling and estimation of population summary statistics, with the following filters: ‘gstacks: --min-mapq 20 --max-clipped 0.2’ and ‘populations: -p 3 -r 0.5’. To avoid the inclusion of paralog loci and poorly mapped regions we excluded RADtags with more than 10 SNPs. The set of used samples is indicated in Table S3.

### Whole genome sequencing

Genomic DNA of 20 Irish mountain hares (details in Table S1) was used to prepare double-indexed individually barcoded libraries, following Meyer and Kircher (2010) with slight modifications (see Giska et al., 2019). These libraries were sequenced in ∼33% of four lanes of an Illumina HiSeq 1500 at CIBIO-InBIO’s New-Gen sequencing platform, to obtain 125 bp paired-end reads. The whole genome sequencing (WGS) data was complemented using similar data from Alpine (*L. t. varronis*, N = 20) and Fennoscandian (*L. t. timidus*, N = 19) specimens analysed previously by Giska et al. (2019). In addition, a dataset of whole genome sequence data at higher individual coverage was put together, including four mountain hare specimens, from Ireland, the Alps, Fennoscandia and the Faroe Islands (6.9-32.8X; from Seixas et al., 2018; Giska et al., 2019; Marques et al., 2020), and five other *Lepus* species: *L. americanus, L. californicus, L. europaeus, L. granatensis*, and *L. castroviejoi* (6.1-21.8X; from Seixas, 2017; Jones et al., 2018; Seixas et al., 2018; Giska et al., 2019) (details in Table S2).

The quality of raw Illumina reads was examined using FastQC (available at https://www.bioinformatics.babraham.ac.uk/projects/fastqc/), and the reads were trimmed for adapters and quality using Trimmomatic, TRAILING:15, SLIDINGWINDOW:4:15, MINLEN:30 (Bolger, Lohse, & Usadel, 2014). Reads were mapped onto the hare pseudo-reference genome, and analyses were repeated using also the Irish mountain hare *de novo* draft reference genome (Marques et al., 2020). PCR duplicates were removed with PICARD (http://broadinstitute.github.io/picard/) and local realignment around indels was done with GATK IndelRealigner.

Complete mitochondrial DNA (mtDNA) genomes were also recovered for each mountain hare individual for which the whole genome was sequenced at lower coverage, using the ABySS assembler (Simpson et al., 2009; Jackman et al., 2017). The standard ABySS run was done with k-mer size of 64 bp (k=64), but modified to k=96 and/or randomly subsampling input reads, when it allowed increasing the contig length. For each individual, the nearly full mtDNA sequence was retrieved by inspecting the longest contigs and then aligning with a mountain hare mtDNA reference (GenBank Acc. Nr KJ397605; Melo-Ferreira, Seixas, Cheng, Mills, & Alves, 2014). Final alignments were produced using ClustalW (Thompson, Higgins, & Gibson, 1994), followed by manual corrections in BioEdit (Hall, 1999).

### Evolutionary relationships, population structure and demography

A Principal Components Analysis (PCA) based on both the mountain hare RAD-seq and WGS datasets was carried out using PLINK 1.9 (Chang et al., 2015) and the single-read sampling approach implemented in ANGSD (Korneliussen, Albrechtsen, & Nielsen, 2014), respectively. In all ANGSD analyses we used the following set of quality filters: -minMapQ 20 -minQ 20, -trim 0, -SNP_pval 1e-6, and genotype likelihoods were estimated using SAMtools model (-GL 1), adjusting additional options to the needs of a particular analysis and data set. For each population in the RAD-seq dataset, we performed the SNP calling using ANGSD applying the following additional options: -doMajorMinor 5, -doMaf 1, -doCounts 1, - setMinDepthInd 4, -doGeno 4, -doPost 1, and assuming the reference variant as ancestral. We removed sites with over 20% of missing data per population, sequencing depth higher than three times the mean depth times the number of samples in each region, and the minor allele frequency was set to remove singletons. Then, using PLINK 1.9 we removed individuals and SNPs with a missing rate higher than 10%, and excluded SNPs failing the Hardy-Weinberg test (p=0.05). For the WGS dataset the following additional filters were applied: -uniqueOnly 1, -remove_bads 1, -baq 2. To minimize linkage, SNPs in both data sets were sampled at least 20 kb apart. To further examine the genetic relationships of mountain hares from the different areas, we used individual pairwise distances to construct a neighbour-joining (NJ) tree based on the whole genome data, and allele counts to construct a TreeMix tree using the whole genome data and RAD-seq data. ANGSD genotype posterior probabilities at 128,019 SNPs filtered for a minimum distance of 20 kb were used to calculate pairwise genetic distances with ngsDist (Vieira, Lassalle, Korneliussen, & Fumagalli, 2016). Branch support values were generated through bootstrapping (100 replicates, 100 SNPs blocks, with replacement), and the trees were inferred using FastME (Lefort, Desper, & Gascuel, 2015) and plotted with FigTree v1.4.4 (available at http://tree.bio.ed.ac.uk/software/figtree/). TreeMix (Pickrell & Pritchard, 2012) analyses (1000 bootstrap replicates of 100 and 20 SNPs blocks for WGS and RAD-seq data, respectively) were performed using allele frequency estimates from ANGSD converted into allele counts at SNPs at least 20 kb apart (136,874 and 1,268 SNPs, for WGS and RAD-seq data, respectively). A high coverage genome of the snowshoe hare was used as outgroup for both the NJ and TreeMix trees.

A mitochondrial DNA phylogeny was inferred using the MultiTypeTree method (Vaughan, Kühnert, Popinga, Welch, & Drummond, 2014) implemented in the Beast v2.6.1 package (Bouckaert et al., 2019). This method is based on a structured coalescent model in which the phylogenetic inference incorporates known structure across samples (here set to Ireland, Alps and Fennoscandia). Three independent replicate runs of 100 million MCMC generations were carried out, using the HKY+G+I site model (determined using jmodeltest2; Darriba, Taboada, Doallo, & Posada, 2012) and a Log Normal relaxed clock (Drummond, Ho, Phillips, & Rambaut, 2006). The runs were assessed for convergence with Tracer v1.7.1 (Rambaut, Drummond, Xie, Baele, & Suchard, 2018), and the first 10% of each run was discarded as burn-in. The resulting trees were merged using LogCombiner and summarized using TreeAnnotator, which is part of the Beast package, and plotted with FigTree v1.4.4. Node ages were converted to years using a mutation rate estimated from the *L. timidus* - *L. granatensis* average corrected mtDNA distance, considering a split time of 1.86 million years (Giska et al., 2019).

Finally, the RAD-seq data was used to model the demographic history of European mountain hares, first considering each sampled area – Ireland, Alps, Fennoscandia – as a separate population and modelled independently, and second using the single-population estimates to guide multi-population models. Using ANGSD, we estimated the site allele frequency likelihoods based on individual genotype likelihoods from bam alignments: -GL 1 -doSaf 1. Following the same procedure applied in the PCA, sites with over 20% of missing data and sequencing depth higher than three times the global depth per population were removed. Individuals with sequencing depth at the particular site lower than four (-setMinDepthInd 4) were also removed. The site frequency spectrum (SFS) for single populations and the 2D SFS were inferred applying the expectation-maximization (EM) algorithm optimization with realSFS program of ANGSD. 2D SFSs were estimated using sites commonly covered across the three populations. SFS for single populations and 2D SFSs were inferred using all sites available after filtering. The coalescent-based method implemented in fastsimcoal v.2.6.0.3 (Excoffier, Dupanloup, Huerta-Sánchez, Sousa, & Foll, 2013) was used to infer the demographic parameters. The separate demographic models for each population were first used to get accurate estimations of current effective population sizes. Then, the demographic history of the divergence of the three considered mountain hare populations was inferred based on two different models (following the neighbour-joining and TreeMix inferences – see Results): 1) considering a simultaneous split (Same Time Model - STM), and 2) considering two splits forward in time, first of the Alpine mountain hares and then between Fennoscandian and Irish mountain hares (Different Time Model - DTM). For the STM and DTM models, the current effective populations sizes were fixed, based on the estimations from the single population models (Table S4), to reduce the large parameter space of the multi-population models. The three individual population scenarios and the two models of population divergence were run for 250,000 simulations and for 100 replicates with random parameter values as starting point. To avoid overfitting of the model, we used a threshold of 10 (-C) for observed SFS entry count, pooling all entries with less than 10 SNPs. We optimized the fit between estimated and observed likelihoods by performing a total of 100 expectation-conditional maximization (ECM) cycles. For the first 50 cycles, the likelihood was computed with monomorphic and polymorphic sites, while for the remaining 50 cycles, we used only polymorphic sites. Parameters were estimated using a maximum composite likelihood from the unfolded SFS, and the run with the best likelihood was retained. The method calculates a maximum observed likelihood by replacing the estimated SFS by the empirical SFS, thus obtaining the maximum possible likelihood. Small differences between the estimated and maximum observed likelihood indicate good fit. For the three-population models, density intervals for parameters values were estimated by parametric bootstrapping using fastsimcoal. The parameters generated from the run with the best likelihood were used to generate 100 pseudo-datasets using the following options: ‘-n 100 -j -d -s0 -x -I -q’. For each of these pseudo-datasets, fastsimcoal was run under the same conditions as the empirical data. In each case, the parameters values were retrieved, and 95% high-density intervals (HDI) were estimated with R using the function *HPDinterval* included in the package *coda* (Plummer, Best, Cowles, & Vines, 2006).

### Genome-wide scans for adaptive differentiation and selective sweeps

Genomic regions underlying local adaptation in a given population are expected to show extraordinary levels of differentiation relative to the other two populations. Using the low individual coverage whole-genome sequencing data, we performed genome scans of the population branch statistics (PBS), which has been shown to have power to detect recent positive selection (Yi et al., 2010). PBS scans were run with PBScan (Hamala & Savolainen, 2019) on genotype likelihoods estimated with ANGSD, using only sites represented by at least six individuals per area. The sites were filtered for minimum coverage of 10 (-setMinDepth), maximum coverage 3 times the average coverage (-setMaxDepth), and the quality filters mentioned earlier. Additionally, we applied a minimum of 4 read counts per base and minor allele frequency maf=0.05. The final data set consisted of 8,978,539 SNPs, including 8,770,557 autosomal SNPs, and the PBScan was run in non-overlapping windows of 50 SNPs. We used Hudson’s F_ST_ for differentiation estimates (-div 0). Outlier detection was based on p-values derived from neutral data simulated with *msms* (Ewing & Hermisson, 2010) using demographic parameters inferred according to the STM and DTM models (Table S4). We simulated 10,000 fragments of 10 kb as the median length of 50 SNPs window in PBS scan was 10,104 bp. We ran PBScan separately with data simulated based on mean demographic parameters of each model, supplied with -ms option. For each underlying demographic model, outliers were defined as genomic regions with at least two consecutive windows with *p* ≤ 0.01. The union of the identified outliers for the two models was retained as candidates for local adaptation. Because the three sampled areas are isolated and thus populations are strongly structured, which could lead to spurious outliers of differentiation, we also used BayPass v.2.2 (Gautier, 2015) to re-assess PBS outliers taking this structure into account. We ran the core model with a contrast statistics (C2; Olazcuaga et al., 2020) based on allele frequencies corrected for structure, using the scaled covariance matrix of population allele frequencies (Ω). The model was run for three possible settings of two groups, the first including the area of interest, and the second containing the two remaining areas. The input file was generated with *poolfstat* R package (Hivert, Leblois, Petit, Gautier, & Vitalis, 2018) from a sync file, and included the same set of autosomal sites as used for PBS. The analysis was run per SNP with default parameters, except -npilot 30 and -d0yij 8. Then, we calculated mean C2 statistics in non-overlapping windows of 50 SNPs, in keeping with window sizes used in the PBS scans. We considered regions with at least two consecutive windows within 0.1% top C2 values as outliers. This procedure was used for three independent runs with different seed numbers, but given highly concordant results across runs we kept the results of a single run.

To validate if the low coverage individual data allowed properly recovering allele frequency differences, we selected 28 SNPs for genotyping in four genomic regions showing increased differentiation in Irish mountain hares (chr03 - *NR3C1*, chr04 - *HMGA2*, chr13 - *DNM3*, chr18 - *EXOC6*). Target SNPs were selected in regions of high differentiation determined using Fisher exact tests implemented in Popoolation2 (Kofler, Pandey, & Schlötterer, 2011). Fisher exact tests were based on sync input files and run with minimum count of four, minimum coverage of ten, maximum coverage 3X the average coverage, non-overlapping windows of 20 kb covered in at least 25%, and the resulting P-values were corrected using the Bonferroni criterion. Six to eight SNPs were genotyped per genomic region in 103 individuals, using the AgenaBio iPLEX technology at Instituto Gulbenkian de Ciência (IGC), Oeiras, Portugal.

We next scanned the genomes of mountain hares for within population signatures of selective sweeps using three statistics: nucleotide diversity (π), Tajima’s D and the composite likelihood ratio (CLR) of SweeD (Pavlidis, Zivkovic, Stamatakis, & Alachiotis, 2013). The de-correlated composite of multiple signals (DCMS) approach (Ma et al., 2015) was used to summarize the statistics. In particular, we tested whether high differentiation coincided with population-specific signatures of selective sweeps, thus allowing differentiating local adaptation from global adaptation (Booker, Yeaman, & Whitlock, 2021). Tajima’s D was calculated in 100 kb windows with 20 kb steps using PoPoolation1 (Kofler, Orozco-terWengel, et al., 2011), based on the mpileup file subsampled to a uniform coverage of 10, the classic option selected (--min-count 1), and the following options: --min- qual 20, --min-covered-fraction 0.25, --dissable-corrections. Nucleotide diversity was calculated using PoPoolation1, with the mpileup file used as input employing the following parameters: --min-count 4, --min-coverage 10, --min-qual 20, --min-covered-fraction 0.25, and maximum coverage 3 times the average coverage. The inference of composite likelihood ratio implemented in SweeD was performed based on ANGSD-derived allele frequency estimates, assuming the European rabbit variant as the ancestral state. Grid size was set to define non-overlapping 20 kb windows along the whole-genome, and CLR values were averaged in 100 kb windows with 20 kb steps. Only windows overlapping across the three inferences were retained for DCMS, which was run using R package MINOTAUR (Verity et al., 2017). Raw statistics were first converted to p-values based on a left-tailed (nucleotide diversity and Tajima’s D) or right-tailed test (SweeD CLR). Then DCMS score was calculated for each window along the genome and visualized as a Manhattan plot with the top 1% genome-wide distribution values indicated. Outliers of DCMS were considered coincidental with the PBS ones if the distance between them was not greater than 80 kb (four windows in the DCMS scan).

### Comparative genomic analyses

To understand the origin of candidate locally adapted variants in the context of the evolution of the genus, we analysed the high coverage whole genome data of six *Lepus* species (1 to 4 individuals per species), including representatives of Irish, Alpine, Fennoscandian and Faroese mountain hares (Table S2). Maximum-likelihood phylogenies were built using RAxML with the GTR+G model (Stamatakis, 2015) for alignments containing all autosomes, and then each genomic region identified as candidate to underlie local adaptation (Table S10). The European rabbit (*Oryctolagus cuniculus*) was used as outgroup in these phylogenetic reconstructions.

We then focused on cases where the mountain hare candidate adaptive variant shared the most recent common ancestor with another species and not with their conspecifics. Such phylogenetic pattern could either result from the retention of ancestral variation or from secondary introgression. Given that introgression would lead to low local sequence divergence to the donor species, we estimated genetic divergence, both raw (*d*_*XY*_) and mutation-scaled divergence (Relative Node Depth (*RND*), using the European rabbit as outgroup) (Feder et al., 2005) in 20 kb overlapping windows with 2 kb step along two chromosomes of interest, 1 and 8, using genomics_general scripts (https://github.com/simonhmartin/genomics_general). Only windows with a minimum of 25% of the sites covered were included for these estimates. To further test whether low empirical local divergence could be simply driven by the variance of the coalescence during species divergence, we simulated the expected local sequence distances under a strict isolation model, following Giska et al. (2019). We first used G-PhoCS (Gronau, Hubisz, Gulko, Danko, & Siepel, 2011) to estimate demographic parameters of the species divergence (effective population sizes, divergence times, and migration rates) using the higher coverage whole-genome data (Table S11). For *L. timidus*-*L. castroviejoi* inferences, G-PhoCS runs were based on 9,256 fragments of 1 kb length sampled along the genome and fulfilling the following criteria: within intergenic regions, at least 50 kb apart, maximum 40% missing sequence after masking repeats with Ns. Estimates of *L. timidus*-*L. europaeus* divergence were from Giska et al. (2019). We then used the estimated demographic parameters in *msms* (Ewing & Hermisson, 2010) to simulate the expected sequence divergence using a mutation rate of 2.8 × 10^−9^ substitutions/site/generation (Giska et al., 2019), and a model with no post-divergence migration. We also simulated an extreme case of a strong selective sweep (selection coefficient s = 0.1) starting 25 kya prior to the species split, forcing the time of allelic divergence between species to coincide with the species split, thus setting a conservative lower bound for the expected divergence under no gene flow.

## Results

### Sequencing and demographic history

The RAD-seq dataset consisted of 62 samples, including 20 from Ireland, 16 samples from the Alps and 26 from Fennoscandia. The whole genomes of 20 Irish mountain hares were sequenced for a total population coverage of 20.5X (mean 1.0X per individual), and combined with similar published mountain hare data from the Alps (N = 20; total coverage 17.1X; mean individual coverage 0.85X) and Fennoscandia (N = 19; total coverage 18.0X; mean individual coverage 0.95X) (Giska et al., 2019). Mitochondrial DNA sequences were reconstructed from the low individual coverage whole genome sequencing data, resulting in an alignment of 16,223 bp (N = 59).

The PCA confirmed the expected strong population structure of the analysed fragmented species (Figure 1B, Figure S1). The Irish mountain hare showed larger differentiation (average pairwise F_ST_ of 0.22 and 0.25, with Fennoscandian and Alpine populations, respectively; F_ST_ = 0.12 between the Alpine and Fennoscandian population; based on WGS data), which agrees with its inferred smaller effective population size (Figure 2), and the distinct mtDNA clade (Figure 1). The genomic neighbour-joining tree suggested a closer relationship between the Irish and Fennoscandian populations (Figure 1C), an evolutionary pattern that was also supported by TreeMix and the mitochondrial DNA haplotype phylogeny (Figure 1D; Figure S2). However, the NJ whole-genome phylogeny showed paraphyly of the Alpine genetic variation (Figure 1C), while the mtDNA phylogeny showed monophyly only in the Irish mountain hare (Figure 1D). Single population demographic models simulated with fastsimcoal showed good fit to the empirical SFSs (difference between the maximum observed and the estimated likelihoods, ΔLhood of 146, 620, and 462 for Irish, Alpine, and Fennoscandian mountain hares, respectively). The estimated effective population sizes were used to infer the history of divergence among the three mountain hare populations using two alternative models (simultaneous split (STM) or a sequential split (DTM)), which showed similar fit to the data (ΔLhood, 3870 and 3802 for the STM and DTM model, respectively, Table S4-C). Similar time-frames for the population split were inferred: 18.4 kya (thousand years ago; assuming 2 years per generation; 95% HDI 17.5 - 22.1 kya) for the STM model, and 21.2 kya (95% HDI 19.1 – 25.3 kya) and 17.8 kya (95% HDI 15.9 – 20.1 kya) for the DTM model (Figure 2; see also Table S4-C for detailed parameter estimates). Slightly different demographic trends were suggested from the STM or DTM models but with overlapping 95% HDI (Figure 2; Table S4-C). Despite the natural uncertainties of molecular calibrations and large confidence intervals of the estimates (Table S4-C), our results are consistent with simultaneous or near-simultaneous fragmentation of the three mountain hare populations occurring at late Pleistocene, possibly around the Last Glacial Maximum.

**Figure 2.**
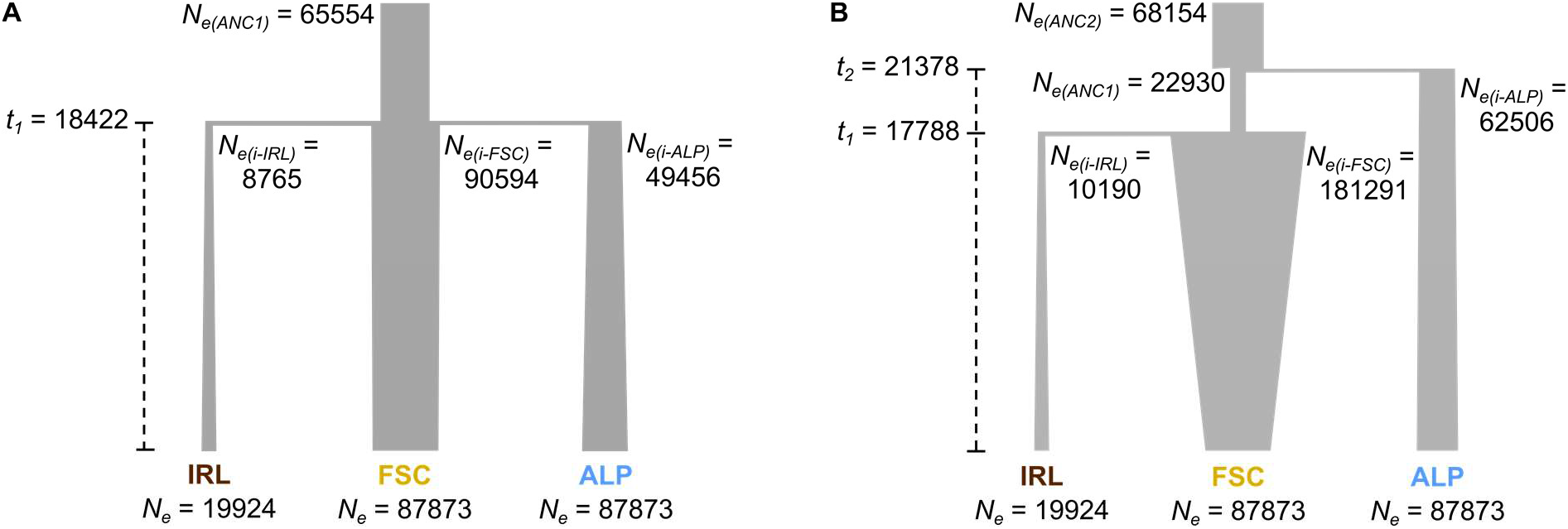
Demographic history of the mountain hare populations. Models were inferred from RAD-seq data. (A) The simultaneous split (STM) model. (B) The sequential split (DTM) model. Parameter estimates are shown: *N*_*e*_ – effective population size in number of diploid individuals (*i* – initial; *ANC* – ancestral); *t* – time of splits in years, considering 2 years per generation (Marboutin & Peroux, 1995). Ireland - IRL, Alps - ALP, Fennoscandia - FSC.

### Differentiation outliers and signatures of selective sweeps

Population branch statistics (PBS) scans between Irish, Alpine and Fennoscandian mountain hares revealed several genomic regions with extreme levels of differentiation. We considered autosomal outlier genomic regions for inter-population differentiation those with at least two consecutive windows of 50 SNPs with PBS of *p* ≤ 0.01 (established from simulations based on the demographic models), confirmed by BayPass contrast scans (Figure S3, Table S5, Table S6). We identified 12 differentiation outliers in Irish, 18 in Alpine, and 9 in Fennoscandian mountain hares. These genomic regions overlapped 14, 9, and 12 protein coding genes, respectively (Figure 3; Table S5). Consistent results were obtained when analyses were based on the *de novo* Irish mountain hare draft genome reference (Figure S4, Table S7, S8). Our independent validation of allele frequency inferences, based on SNP genotyping in four candidate regions in the Irish mountain hare, confirmed the patterns inferred from the whole genome data of low individual coverage (Figure S7, Table S9).

**Figure 3.**
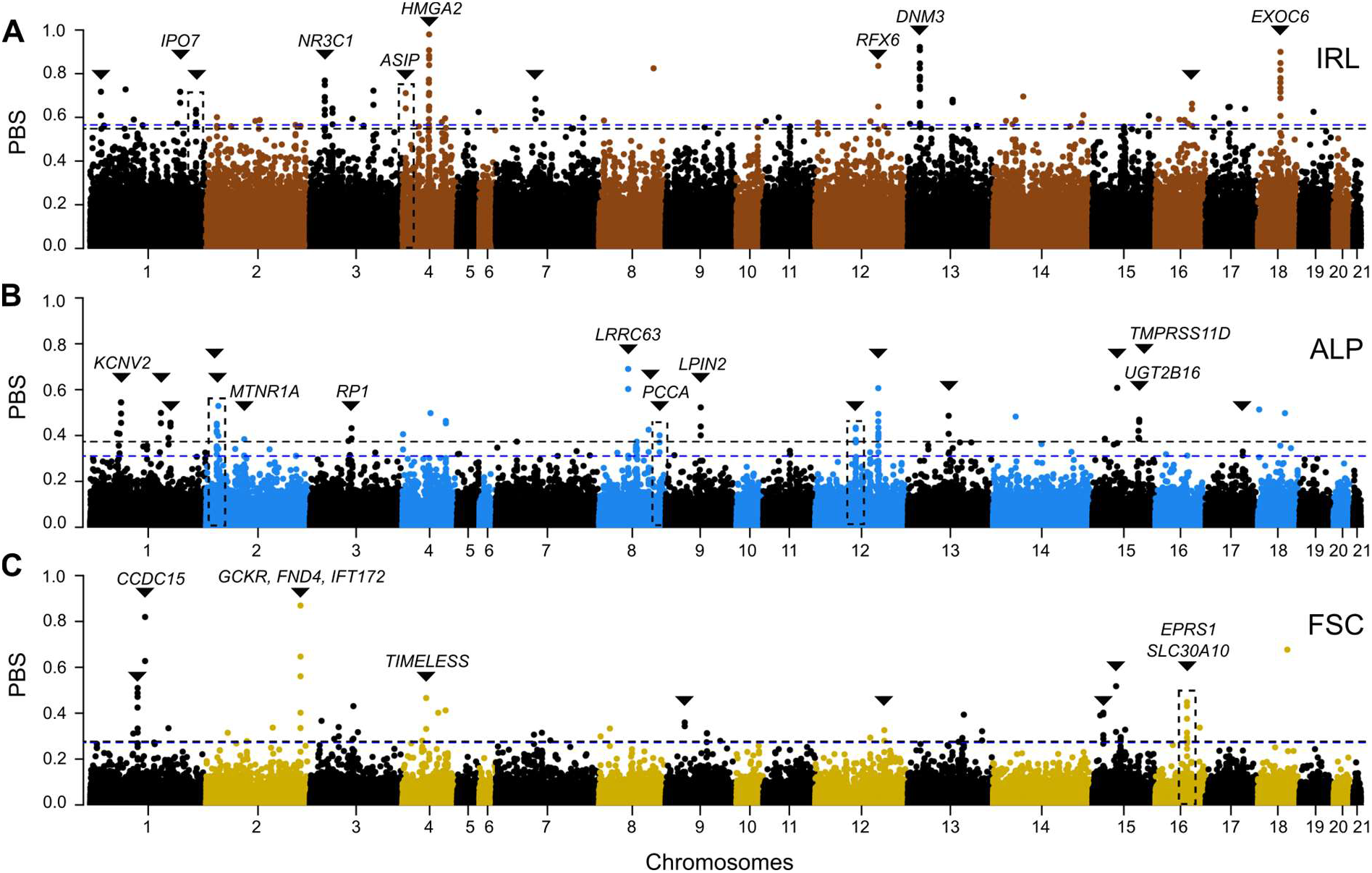
Population branch statistic and selective sweep (DCMS) scans along the autosomes for each mountain hare population (A – Ireland; B – Alps; C – Fennoscandia). The peaks marked with black triangles represent regions with at least two consecutive windows with PBS p ≤ 0.01 (blue dashed horizontal lines indicate thresholds based on the STM model simulations, and black dashed horizontal lines indicate thresholds based on the DTM model simulation), confirmed by BayPass contrast analysis (see Figure S3). Codes indicate genes within the outlier regions (see Table S5). Dashed rectangles indicate PBS peaks that overlap with signatures of population-specific selective sweeps, evidenced by DCMS analysis (see Figure S6 and Table S10). See Figure S5 for PBS scans along chromosome X.

The combined evidence of PBS and BayPass, identifying differentiation outliers, and population-specific DCMS, identifying intra-population selective sweeps (Figure S6, Table S10) pinpointed two candidate genomic regions in Irish mountain hares, one overlapping gene *ASIP*, three candidates in Alpine mountain hares, one overlapping gene *PCCA*, and two candidates in Fennoscandian mountain hares, one overlapping genes *EPRS1 and SLC30A10*. Additional selective sweep signals were detected in other PBS peaks, but were found to be shared among two or all three populations, suggesting possible global and not local selection (Table S10). These regions under global selection included genes *IPO7, GCKR, FNDC4, IFT172, HMGA2, RFX6, DNM3, and EXOC6*.

### Comparative genomics

From the analyses of the higher coverage whole genome data across species, we found that local gene trees of candidate regions for local adaptation predominantly recovered the mountain hare as monophyletic lineage (Figure 4). However, in two cases, the local phylogenetic tree was discordant from the presumable species tree, with the candidate variant grouping with variants from another species with strong bootstrap support (Figure 4C,E). These patterns of shared variation between species could either result from interspecies incomplete lineage sorting or secondary introgression. We found that within both these regions, chromosome 1 (177.8 Mb, unknown gene content) and chromosome 8 (101.7 Mb, *PCCA*), the divergence to the potential donor species is lower than intraspecific divergence between populations, and the pattern is not influenced by local mutation rate when controlled using the *RND* (Figure 5 A,C). Simulating the expected distribution of divergence between species showed that, in both cases, the empirical divergence between the mountain hare and the presumable donor species along the candidate region is smaller than 1% lowest values simulated under a species divergence model with strong selection and no post-split gene flow (Figure 5). In the case of the candidate region on chromosome 8, this pattern was found along a predominant portion of the region, while for the chromosome 1 it affected a shorter fragment of the candidate genomic stretch. These results suggest that in both cases of phylogenetic discordance, introgressive hybridization introduced the adaptive variants in the mountain hare, from *L. europaeus* into Irish *L. timidus* (unknown gene content) and from *L. castroviejoi* into Alpine *L. timidus* (gene *PCCA*) (Figure 4).

**Figure 4.**
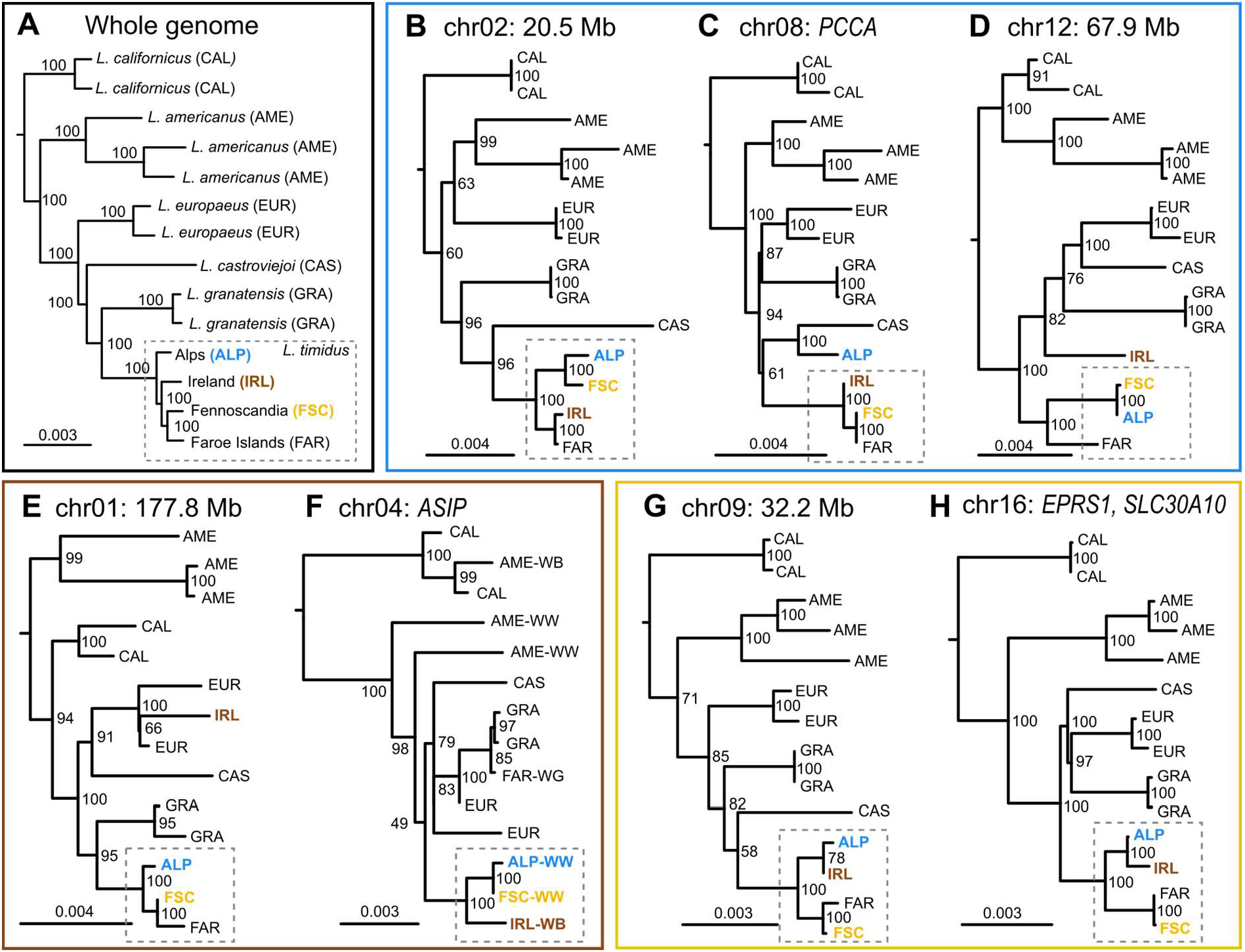
Maximum-likelihood trees for the whole genome and local trees of candidate regions for local adaptation in the mountain hare (genes in the regions are noted, unless no annotation is known, in which case the coordinate is noted). (A) Global tree based on the concatenated whole genome. Local phylogenies of candidate regions in (B-D) Alpine mountain hares, (E-F) Irish mountain hares, and (G-H) Fennoscandian mountain hares. In the *ASIP* phylogeny, notations WW (winter-white), WB (winter-brown) and WG (winter-grey), indicate winter colouration morphs of the sequenced individual.

**Figure 5.**
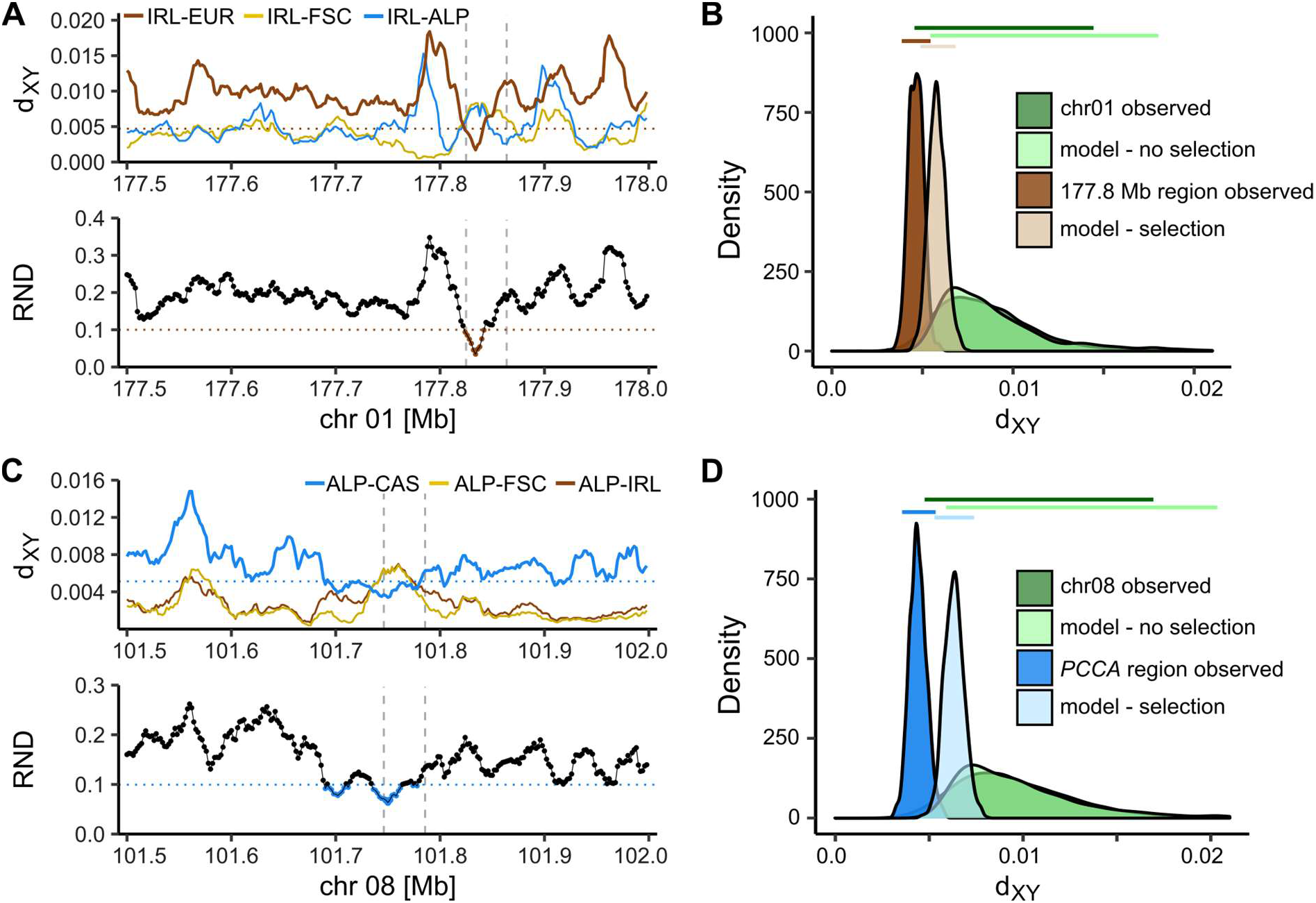
Signatures of introgression. Left panels: scan of *d*_*XY*_ (top; horizontal dotted line indicates 1% lowest values in interspecific *d*_*XY*_ simulated under a model with selection; IRL - Ireland, ALP - Alps, FSC - Fennoscandia, EUR - *L. europaeus*, CAS - *L. castroviejoi*) and *RND* between focal mountain hare and potential donor species (bottom; horizontal dotted line indicates 1% lowest values in the observed chromosome-wide distribution), with vertical dashed lines representing the candidate regions. Right panels: distribution of *d*_*XY*_ between the focal mountain hare population and the putative donor species, including the observed distribution along the whole chromosome and the candidate region, and simulated distribution under the model with and without selection. Top horizontal bars represent 95% range for each distribution. (A-B) Irish *L. timidus* - candidate region chr01:177.8 Mb and possible introgression from *L. europaeus*, EUR. (C-D) Alpine *L. timidus* - candidate region chr08:101.7 Mb, overlapping *PCCA* gene, and possible introgression from *L. castroviejoi*, CAS.

## Discussion

In the present work we studied genomic variation of a typically cold-adapted species, currently fragmented into isolated populations in Western Europe, characterized by vast environmental differences. We identified candidate targets of positive selection, the biologically relevant genetic variation underlying local adaptation, and the evolutionary origin of some of these variants through introgression from related species.

Our demographic modelling suggested that the current Irish, Alpine and Fennoscandian mountain hare populations resulted from a nearly simultaneous fragmentation from a common gene pool in Europe during the late Pleistocene, possibly around the Last Glacial Maximum ∼20 kya (Figure 2; Table S4). The inferred split event appears compatible with a phylogeographic pattern characterised by periodical shifts in geographical range distribution accompanied by further fragmentation caused by retreats and advances of the ice streams during deglaciation, typical of cold-adapted species (Stewart, Lister, Barnes, & Dalen, 2010). Indeed, previous analyses of mountain hares, based on few markers, have shown variation sharing across the range of the species, which is compatible with recent fragmentation (Hamill et al., 2006; Melo-Ferreira et al., 2007; Smith et al., 2017). This, together with our results, fits a scenario in which a single mountain hare population became fragmented during glacial retreat. Our inference sets the upper limit for the time-frame of local adaptation in the species in the newly colonized territories at late Pleistocene.

Our scans of population branch statistics identified highly differentiated regions of the genome in all three analysed isolated populations (Figure 3, Table S5). These extreme differentiation regions could be spurious signals of extreme differentiation, given the structured nature of the contrasted populations (Hoban et al., 2016), but our analyses showed that the patterns remain when taking the covariance of allele frequencies into account, thus controlling for population structure (Figure S3, Table S6) (Gautier, 2015). Thus, spurious patterns of variation resembling local adaptation could result from spatially uniform positive or negative selection, decreasing variation at linked sites, and increasing the variance of pairwise differentiation estimated locally in the genome (Cruickshank & Hahn, 2014; Hoban et al., 2016). While our candidate PBS outliers were population-specific, we did detect several instances of candidate regions where a signal of selective sweep based on intra-population statistics was shared across two or three populations analysed, which could result from simultaneous sweeps or uniform background selection (Figure S6, Table S10). Nevertheless, we identified seven genomic regions with population-specific signatures of extreme local differentiation and population-specific selective sweeps.

We inspected the gene content of these candidates for local adaptation and found that some gene functions match phenotypic uniqueness of the populations. For instance, in the Irish mountain hare, the pigmentation gene *ASIP* was identified as a candidate. This protein is an antagonist of the melanocortin 1 receptor (MC1R), present in the membranes of melanocytes, shifting the production of pigment from dark eumelanin to light phaeomelanin or inhibiting the production of pigment (Hoekstra, 2006). It is a well-known pigmentation gene that underlies adaptive coat colour variation in many mammals (Kerns et al., 2004; Fontanesi et al., 2010; Barrett et al., 2019; Miranda et al., 2021). Mountain hares typically undergo brown-white seasonal moults to remain camouflaged year-round on seasonal snow (Zimova et al., 2018). Yet, winter coats in Irish mountain hares present typically a unique brown winter coat pattern. This suggests a local adaptation, as individuals with white coat would have higher fitness costs by coat colour mismatch due to absence of significant or prolonged snowfall in Ireland (Zimova, Mills, & Nowak, 2016). *ASIP* has been previously linked with coat colour variation in hares, namely the winter brown morphs of snowshoe hares (*L. americanus*) in the Pacific Northwest of the USA (Jones et al., 2018) and the winter grey morph of Faroese mountain hares (Giska et al., 2019) via *cis*-regulatory changes. Our results thus suggest that *ASIP* is also a strong candidate to be involved in the determination of the winter brown colour morph of the Irish mountain hare. In Alpine mountain hares, which inhabit high altitude environments up to ∼4000 m above sea level (Hackländer & Jenny, 2011; Angerbjorn & Schai-Braun, 2021), the *PCCA* gene showed pattern of population specific positive selection. This gene encodes the propionyl-CoA carboxylase enzyme responsible for catabolism of odd chain fatty acids (OCFAs) by carboxylation of their metabolite, propionyl-CoA. It was shown that due to cold exposure the level of OCFAs in the white adipose tissue of mice got changed (Xu et al., 2019). Propionyl-CoA can be transported from mitochondria to the cytoplasm and there used as an intermediate in the Krebs cycle to fight hypothermia when mitochondrial energy production is weakened. It has been hypothesized that OCFAs may increase energy production when temperature is low. Thus, *PCCA* may be involved in adaptations to cold Alpine habitat by intensifying the use of OCFAs as energy source. In Fennoscandian mountain hares, we found evidence of selection overlapping the *SLC30A10* and *EPRS1* genes, which in human GWAS studies have been linked to body mass, bone density and adipose tissue physiology (Arif et al., 2017), and may thus underlie larger body sizes of Fennoscandian hares. Among the genes with signals of global selection (i.e. overlapping signs of selective sweeps in several populations) (Figure S6, Table S10), *HMGA2* has been vastly linked to body size differences in mammals, including dwarfism in rabbits (Jones et al., 2008; Carneiro et al., 2017), and may underlie body size differences across mountain hare populations. The functional relevance of the inferred adaptive variation remains naturally tentative (see complete list of candidate genes in Table S5), but our results can guide future functional testing to validate these hypotheses.

We then analysed the candidate adaptive genetic variation of the mountain hare in the context of the evolutionary history across species. When comparing local and whole-genome phylogenies, we found that the local phylogenies of most of the candidate genomic regions, recovered the mountain hare as monophyletic (Figure 4). This suggests that most of the adaptive variation inferred here is species-specific. With the current dataset, we cannot determine whether adaptation resulted from standing variation or from *de novo* mutation, but standing variation appears more plausible, given the recent post-glacial time-frame of the divergence (Barrett & Schluter, 2008) (Figure 2). However, in some cases we found that the local phylogenetic signal was discordant from the species-tree, with the candidate variant grouping with another species. Our simulations show that for *PCCA*, the empirical divergence between the broom hare found in the Cantabrian mountains in northern Spain and the Alpine mountain hare variant is lower than expected under a no-gene flow scenario, and hence originated from introgression (Figures 4 and 5). These species have been shown to be involved in ancient hybridization (Melo-Ferreira et al., 2012) and this introgression may therefore witness those ancient events. We also found evidence of likely ancient introgression from the European hare into the Irish mountain hare (Figures 4 and 5), but in this case no known gene was annotated to this genomic region, and the source of adaptive variation remains elusive. Even though our approach was rather conservative, these results confirm that adaptive variants may originate from variation introduced through introgression. This can be particularly common in hares, given both historical and contemporary contact between species resulting in interspecific exchanges along the diversification of the genus (Ferreira et al., 2021).

While most of the candidate genes revealed here had not been implicated in known instances of adaptive evolution in hares, the presence of the agouti signalling protein gene (*ASIP*) among candidates for local adaptation appears recurrent (Jones et al., 2018; Giska et al., 2019; this work). In both snowshoe hares and Faroese mountain hares, the variant causing non-white winter morphs introgressed from closely related species, i.e. from black tailed jackrabbits (*L. californicus*) and Iberian hares (*L. granatensis*), respectively. Here we found no evidence of introgression of the Irish mountain hare winter-brown variant (Figure 4), which appears distinct from that generating the grey morph in Faroese mountain hares (Figure S8), and suggests adaptation from species-specific standing genetic variation. In the absence of admixture mapping in natural populations (which is difficult given the isolated nature of the concerned populations) or experimental crosses, the suggestion that *ASIP* underlies the winter brown colour morph in Irish mountain hares remains tentative. However, it is striking that diverse evolutionary pathways recruiting the same gene appear to recurrently promote local adaptation in independent hare populations or species, reinforcing the role of this gene as an evolutionary hotspot for adaptive variation in mammals.

Species remain the central units for conservation assessment and management in national and regional plans, including for the International Union of Conservation of Nature (IUCN) Red Lists and action plans. Yet, this emphasis on species for conservation is controversial, because it fails to consider the continuum of genetic variation, from populations to species (Coates, Byrne, & Moritz, 2018). Moreover, it also fails to consider local adaptation and the unique genetic variation underlying adaptive traits (Teixeira & Huber, 2021). Our work shows that despite the recent common ancestry of the analysed populations of the mountain hare, likely dating back to the Last Glacial Maximum, each of the local populations contains genetic variation that is not present elsewhere. Furthermore, we show that part of that variation, which may be much larger than our conservative approach detected, underlies adaptation to local conditions. This unique adaptive genetic variation is therefore essential for the survival of these organisms in the local environments, where they are cornerstones of the ecosystems, making simplistic species-wide conservation actions, such as unsupervised translocations, ineffective. Yet, identifying genetic variants within species that may seed adaptive responses to climate change, such as those underling winter brown coats that will likely be favoured given projected snow cover decreases (Mills et al., 2018), may also guide the introduction of standing adaptive variation upon which selection can act. As population genomic studies become common, detecting intraspecific adaptive variation will provide information that needs to be incorporated efficiently into conservation planning and practice, fully integrating the evolutionary process in conservation (Mills et al., 2018; Hoelzel, Bruford, & Fleischer, 2019). Regardless of the taxonomic status of the studied mountain hare populations, which are classified as subspecies (Angerbjorn & Schai-Braun, 2021), our work underscores that conservation efforts should take the adaptive uniqueness of the endemisms into account. Such evidences should be considered when assessing species conservation status in the scope of Union of Conservation of Nature (IUCN) Red Lists and action plans.

## Acknowledgments

This work was supported by Fundação para a Ciência e a Tecnologia (FCT) (project grant “2CHANGE” – PTDC/BIA-EVL/28124/2017, co-funded by the European Regional Development Fund through COMPETE 2020, POCI-01-0145-FEDER-028124). J.M.-F. was supported by an FCT CEEC contract (CEECIND/00372/2018). W.I.M, P.A.P, and N.R were supported by the Environment & Heritage Service and Queen’s University Belfast through the Quercus partnership and by the National Parks and Wildlife Service. Instrumentation, laboratory, and computational support was provided by CIBIO NEWGEN sequencing platform, supported by the European Union’s Seventh Framework Program for research, technological development and demonstration under grant agreement no. 286431, and the Montpellier Bioinformatics Biodiversity platform supported by the LabEx CeMEB, an ANR “Investissements d’avenir” program (ANR-10-LABX-04-01). Additional support was obtained from COMPETE2020, PORTUGAL2020 and ERDF (POCI-01-0145-FEDER-022184). All sampled used in this work were collected before the Nagoya protocol came into force where applicable. Most samples were kindly donated by hunters during the regular permitted hunting season or were kindly provided by researchers. We thank F. Suchentrunk, A. Angerbjörn, E. Randi, J. Letty, and Z. Boratynski for kindly providing some of the samples used in this work. Irish mountain hare genetic material was collected under a licence to take animals for scientific purposes issued by the Wildlife Licensing Unit, National Parks & Wildlife Service (NPWS), Department of Housing, Local Government and Heritage. We thank the Irish Coursing Club (ICC) for providing access to hares, and in particular Mr D.J. Histon, Chief Executive Officer, and the Borris-in-Ossory coursing club, for supporting the research. This work has been developed in the framework of the Laboratoire International Associé (LIA) “Biodiversity and Evolution” supported by InEE (CNRS, France) and FCT (Portugal).

## Data Archiving

The raw sequencing data will be deposited in the NCBI Sequence Read Archive (SRA) and the complete mitochondrial DNA sequences in GenBank.

## Author Contributions

J.M.-F. coordinated the study; I.G., J.M.-F., P.C.A., P.A.P., W.I.M., and N.R. designed research; P.A.P., W.I.M., N.R., H.J., and K.H. contributed with sampling; L.F. generated data; J.M.-F. and P.B. provided computational resources and support; I.G., J.P. and J.M.-F. analysed data; J.M.-F. and I.G. wrote the paper with contribution of J.P.; all authors interpreted the results and read, revised, and approved the final version of the manuscript.

