## Supplementary material for "The evolutionary pathways for local adaptation in mountain hares": Figure S

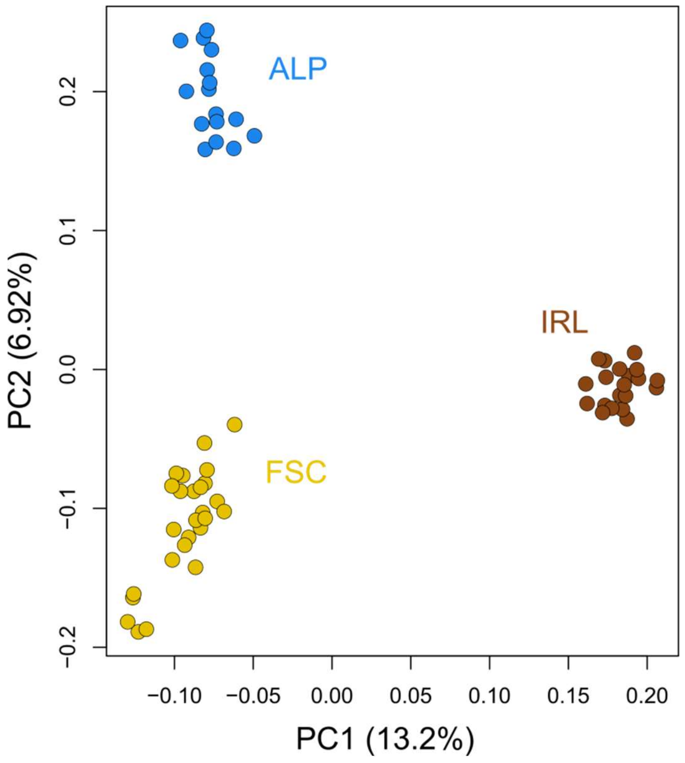


**Figure S1.** Population structure: Principal Components Analysis (PCA) with the proportion of variance explained by the first two components. PCA based on the RAD-seq data, including 1,196 SNPs after filtering for missing data (max. 10% per SNP and individual), Hardy-Weinberg equilibrium and sampling SNPs for a minimum of 20 kb distance; constructed using PLINK and SNPs called with ANGSD.

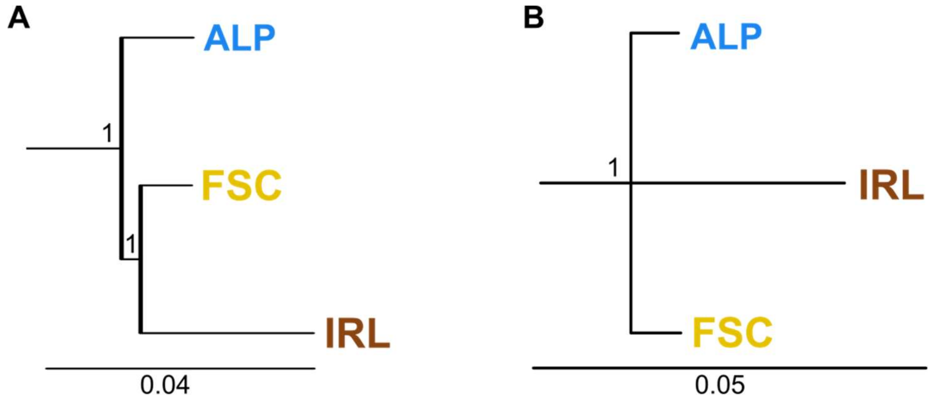


**Figure S2.** Population TreeMix tree based on allele frequencies, with bootstrap support values derived from 1,000 bootstrap replicates and the snowshoe hare (*Lepus americanus*) used as outgroup. **(A)** Tree based on the WGS low individual coverage data, constructed using 136,874 polymorphic sites sampled at least 20 kb apart along the genome. **(B)** Tree based on the RAD-seq data using 1,268 polymorphic sites sampled at least 20 kb apart.

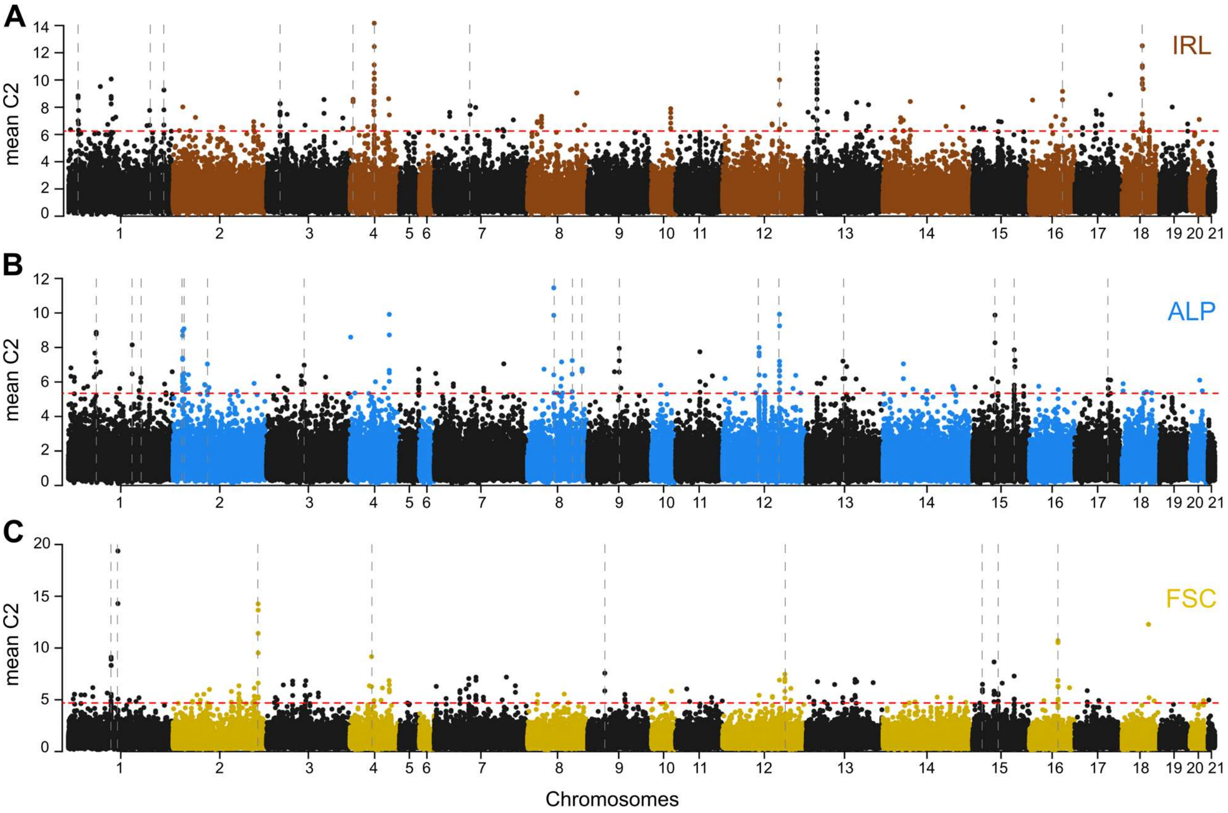


**Figure S3.** Manhattan plots of BayPass contrast scans. The C2 contrast values are plotted as means per non-overlapping windows of 50 SNPs along the chromosomes. The horizontal dashed red line represents 0.1% top autosomal windows based on the observed data set. The vertical grey dashed lines represent the final candidate regions of local adaptation overlapping between PBS and BayPass scans. **(A)** Ireland. **(B)** Alps. **(C)** Fennoscandia.

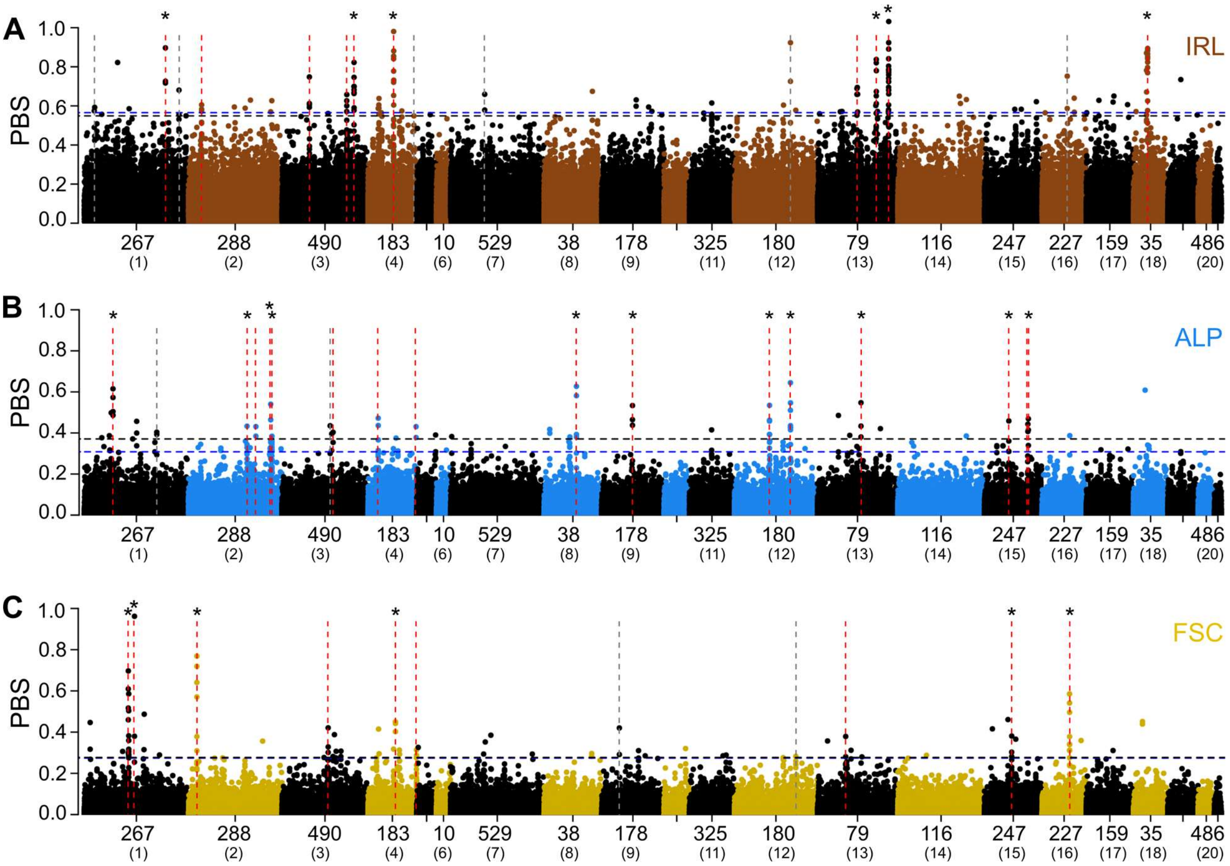


**Figure S4.** Manhattan plots of PBS scan when using data mapped to the mountain hare reference. Horizontal dashed lines represent p=0.01 thresholds based on the simulations of the two demographic models: blue - STM model and black - DTM model. Vertical dashed lines represent PBS outlier regions, confirmed also by BayPass C2 contrast (see Tables S5-S8): red - for data mapped to the mountain hare reference, grey - only for data mapped to the hare pseudo-reference. Regions indicated by asterisks overlap between the two references. The main x-axis label represents scaffolds of the mountain hare reference while the corresponding chromosomes of the hare pseudo-reference are below in parentheses (based on the rabbit OryCun2.0 genome). **(A)** Ireland. **(B)** Alps. **(C)** Fennoscandia.

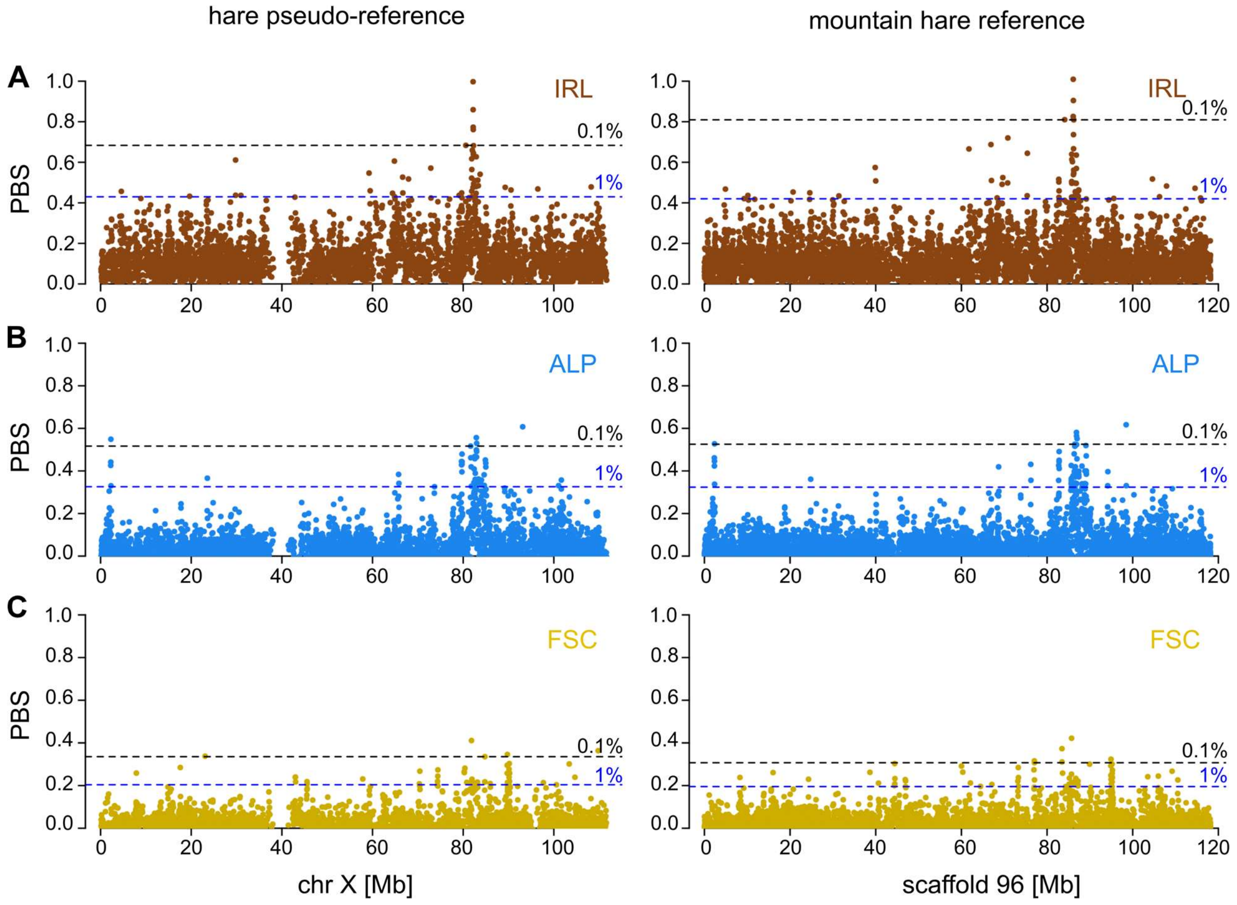


**Figure S5.** Distribution of PBS values along chromosome X. Left panel shows PBS scan for data mapped to the hare pseudo-reference, while right panel shows PBS scan for data mapped to the mountain hare reference. The dashed lines represent top 0.1% and 1% values along the chromosome. **(A)** Ireland. **(B)** Alps. **(C)** Fennoscandia.

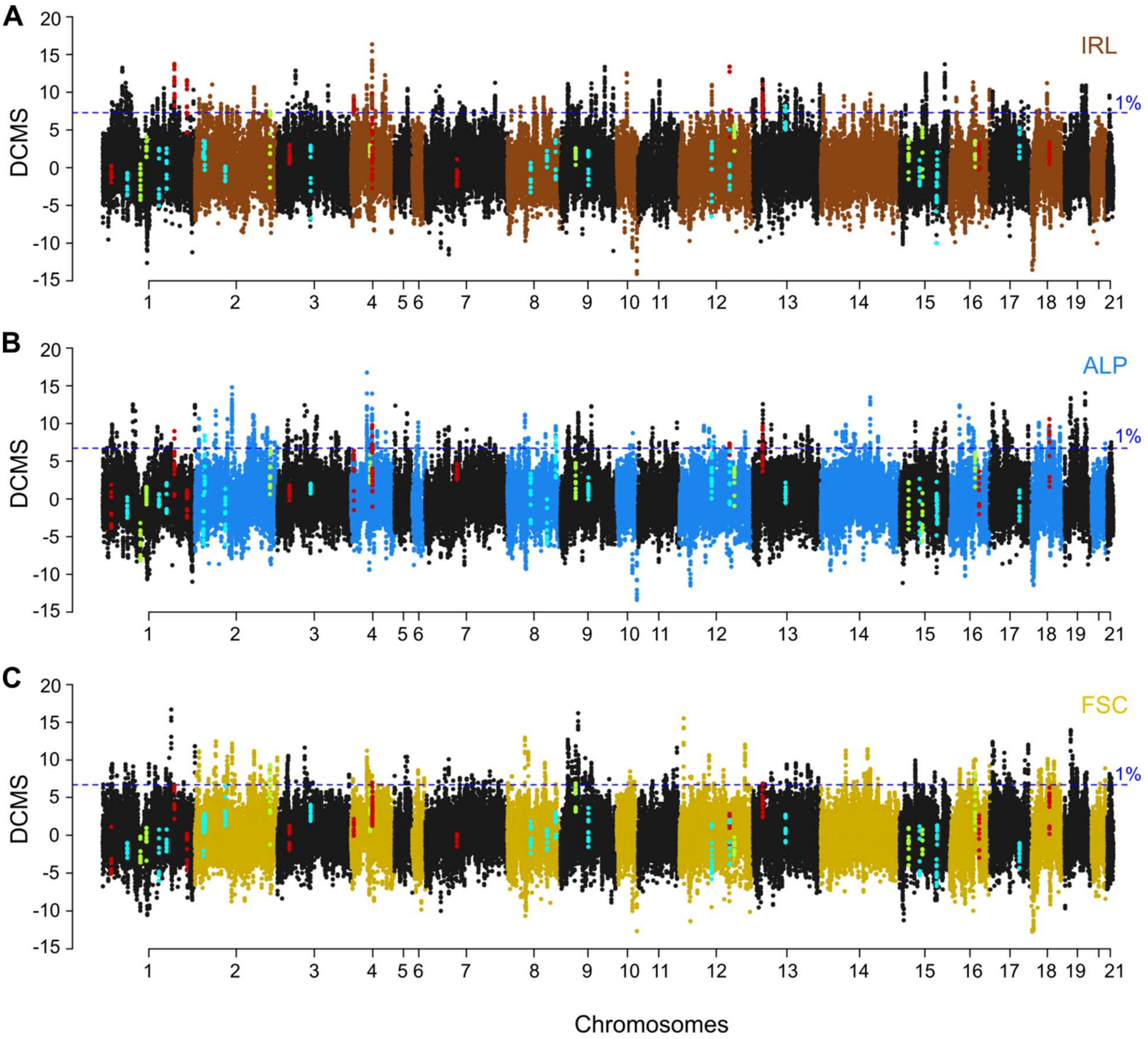


**Figure S6.** De-correlated composite of multiple signals (DCMS) based on three intra-population statistics: SweeD Composite Likelihood Ratio (CLR), Tajima's D and nucleotide diversity (π). The highlighted windows indicate PBS outlier regions extended by three 20kb windows on each side: red - IRL, cyan - ALP, lime - FSC (see Table S5). The horizontal blue line represents top 1% values of autosomal DCMS score distribution in each population. **(A)** Ireland. **(B)** Alps. **(C)** Fennoscandia.

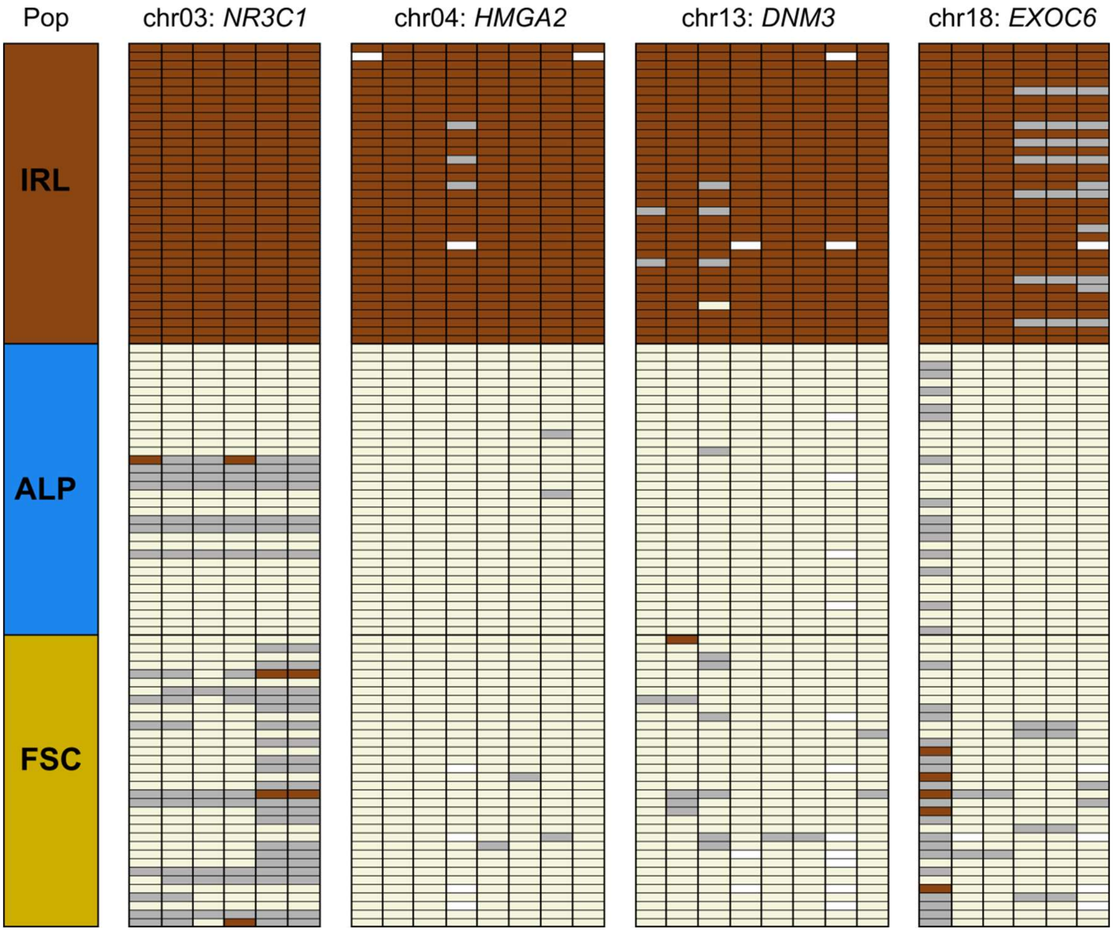


**Figure S7.** Genotypes at 28 loci spanning four PBS outlier regions in Irish hares. Rows represent specimens and columns represent genotyped loci. Brown - homozygous for the Irish variant, beige - homozygous for the alternative variant, gray - heterozygous, white - missing data, IRL - Ireland, ALP - Alps, FSC - Fennoscandia; see Table S5 for details on the outlier regions and Table S9 for details on the genotypes.

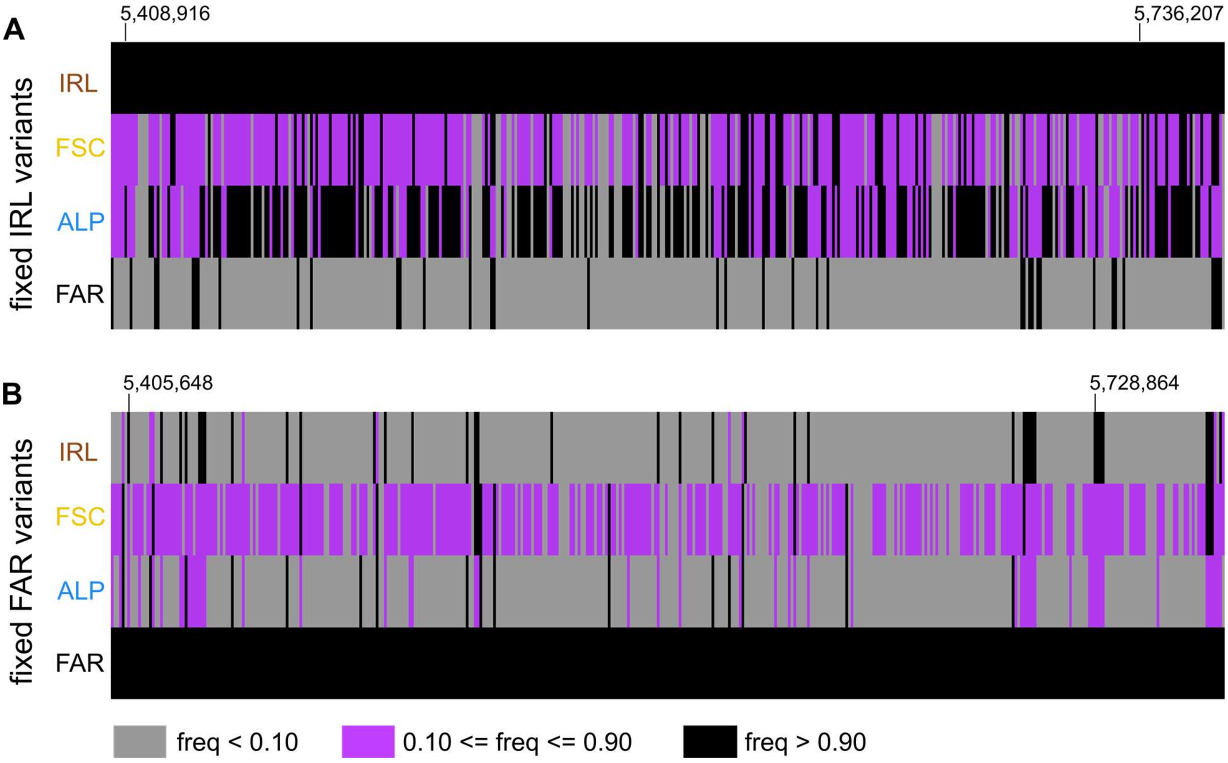


**Figure S8.** Population allele frequencies along the *ASIP* region. Coordinates [bp] are indicated above the Irish panels; IRL - Ireland, ALP - Alps, FSC - Fennoscandia, FAR - Faroe Islands. **(A)** Variants fixed in Irish population. **(B)** Variants fixed in Faroese population.
